# Rational Design of Frontline Institutional Phage Cocktail for the Treatment of Nosocomial *Enterobacter cloacae* Complex Infections

**DOI:** 10.1101/2024.06.30.601436

**Authors:** Dinesh Subedi, Fernando Gordillo Altamirano, Rylee Deehan, Avindya Perera, Ruzeen Patwa, Xenia Kostoulias, Denis Korneev, Luke Blakeway, Nenad Macesic, Anton Y Peleg, Jeremy J Barr

## Abstract

Phage therapy is a promising strategy to treat antimicrobial-resistant infections. Currently, phage therapy applications span personalised treatments that are tailored for a given patient’s infection, through to the use of pre-established cocktails of virulent phages against clinically relevant pathogens. However, both approaches face challenges, with personalised phage therapy being time-consuming and requiring a phage match to a patient’s infection, while phage cocktails may not be effective against a patient’s specific strain. The Alfred Hospital in Melbourne, Australia has reported an ongoing outbreak of infections by the *Enterobacter cloacae* complex (ECC), a group of emerging multidrug-resistant pathogens responsible for considerable morbidity and mortality. Utilising the hospital’s strain collection, built over the last decade, we established an initial three-phage product with 54% ECC coverage that effectively reduced bacterial loads (>99%) in septicaemic mice. We then iteratively improved this product by enhancing phage killing efficiency using phage training and expanded host range through targeted phage isolation against low-coverage ECC strains. This iterative optimisation led to the creation of the product *Entelli-02*, containing five well characterised virulent phages that target clinical ECC strains through distinct bacterial cell surface receptors. Importantly, *Entelli-02* exhibits broad host coverage (99%) and efficacy (92%) against The Alfred Hospital’s ECC strain collection (*n* = 156). We produced this as a therapeutic-grade product, verified and endotoxin unit compliant, ready for use. This approach integrated academic phage research with clinical insights to produce the phage product *Entelli-02* as an institution-specific phage cocktail with frontline efficacy and on-demand availability.

**SUMMARY:** *In brief:* We developed a phage product containing five phages with frontline potential to address infections caused by multidrug-resistant *Enterobacter cloacae* complex.

## Introduction

The emergence and spread of antimicrobial resistance (AMR) poses a serious threat to global health and calls for alternative strategies to combat bacterial infections^1^. Phage therapy, which involves the use of virulent bacteriophages (phages) that can infect and destroy bacteria, has garnered renewed attention as one potential solution to the AMR crisis^2^. Phage therapy offers several advantages over conventional antibiotics, such as high specificity, self-replication, low toxicity, and adaptability to changing bacterial pathogens^3^. Combinations of multiple phages, which are colloquially known as phage cocktails, are often used to broaden the antimicrobial spectrum. Phage cocktails can be produced *a priori* against a target pathogen or group of bacteria^4^. Despite these benefits, clinical trials employing phage cocktails have shown limited clinical efficacy^5–12^. Treatment failure was attributed to target strain divergence, phage stability issues, and the mismatch between phage infectivity on the bacterial isolates the cocktails were constructed upon versus the clinical strains that were eventually treated. In contrast, personalised phage therapy involves the identification and use of a phage with demonstrated activity against a patient’s bacterial infection^13,14^. This approach is often followed by adapting or training the phages to improve their effectiveness, small-scale production, and bespoke treatment. While personalised phage therapy approaches have shown promising results with >70% treatment efficacy^15^, their application presents the additional complexity of requiring rapid phage characterisation, safety and efficacy testing, and administration of the product, all within a practical timeline. A combination of both approaches, with the initial use of a predefined broad-spectrum phage cocktail, while transitioning to personalised approaches as needed, is an appealing strategy for rapid treatment of antibiotic-resistant bacterial infections.

To bridge the divide between broad-spectrum phage cocktails and personalised therapy, we designed a phage product that was targeted towards a high-risk, multi-clonal outbreak of a nosocomial pathogen at a local hospital. We targeted *Enterobacter cloacae* complex (ECC), which is an emerging group of nosocomial pathogens that pose a significant threat to human health due to their acquisition of virulence and AMR determinants. Classified among the ESKAPE pathogens, this complex encompasses clinically relevant species such as *E. cloacae, E. asburiae, E. hormaechei, E. kobei*, and *E. ludwigii,* which can cause diverse infections, such as pneumonia, urinary tract infections, intraabdominal infection, and bacteraemia^16,17^. Notably, these pathogens have considerable epidemic potential, having contributed to a global surge in carbapenem-resistant and extended-spectrum beta-lactamase-producing phenotypes^18^, and were responsible for numerous clonal hospital outbreaks with limited treatment options^19–21^. As a result, ECC is increasingly associated with severe AMR infections in health care settings and was associated with >200,000 deaths globally in 2019 alone^22–27^.

The Alfred Hospital, a tertiary referral centre in Melbourne, Australia, has reported an ongoing outbreak of carbapenemase-producing *Enterobacterales* infections over the last decade^23,28,29^. A retrospective analysis of bloodstream infections at this centre noted ECC contributed ∼20 cases per year, with a recent surge in ECC infections from 2018-2021^29^. Given this clinical burden, there is an urgent need to develop alternative treatments against this endemic, nosocomial, AMR pathogen. To address this, we developed a tailored phage product that is suitable for frontline use against this ECC outbreak at The Alfred Hospital. We developed a standardised approach that combines academic phage research with clinical insights that was backed by an extensive collection of 206 clinical ECC strains collected over the past decade. This enabled the creation of *Entelli-02*, an institution-specific phage cocktail that not only demonstrates frontline efficacy but ensures rapid availability of an effective antimicrobial. Our approach bridges personalised phage therapy with a broad-spectrum phage product that is tailored toward a given hospital’s (*i.e.,* institution) pathogen profile and addresses the urgent need for effective treatment options against AMR pathogens in healthcare settings.

## Results

### Isolation and characterisation of phages against *Enterobacter* isolates from The Alfred Hospital

The Alfred Hospital in Melbourne, Australia, has reported an ongoing outbreak of nosocomial infections with limited treatment options and high mortality rates^29^. Of particular concern is the *Enterobacter cloacae* complex (ECC), which is an emerging AMR threat with considerable epidemic potential^22^. Here, we explored the potential for phages as a frontline antimicrobial solution for The Alfred’s ECC outbreak. Beginning with 206 clinical ECC isolates collected over a 10-year period, we selected a subset of 36 ECC isolates, that represent the sequence type (ST) diversity of the full strain collection (Supplementary data 1). We then constructed an initial phage library against these isolates. After isolating and purifying 21 novel phages, we screened their plaque-forming capacity against our strain subset to obtain a host range map (Fig 1A). Of the 36 isolates, 34 were susceptible to at least one phage. However, considering the time and cost associated with characterising and producing each therapeutic phage, we aimed to build a phage combination that offers the broadest host range with the fewest possible phages combinations. Based on the complementary broad-spectrum host range, we selected øEnA02, øEnC07 and øEnC15 to produce the initial ECC phage combination (cocktail-V1), which provided 61% coverage against the 36 sub-selected ECC hosts (Fig 1A). All phages in the cocktail were sequenced and their genomes were analysed to ensure they did not carry genes known to allow lysogeny or virulence, including integrases, recombinase, mobile genetic elements, or genes encoding antibiotic resistance or toxicity. Based on genome similarity, we classified the phages at the genus level (Supplementary Fig 1 and 2, Fig 1B). Electron microscopy imaging (Fig 1C-E) revealed icosahedral capsids and sheathed contractile tails with lateral tail fibres, consistent with the Tevenvirinae subfamily within the *Straboviridae* family^30^. Importantly, therapeutically relevant bacteriophages must effectively supress target strains and be capable of efficient replication. We evaluated the replicative characteristics of the three phages on their host of isolation using the one-step growth curve, revealing that øEnA02 had a burst size of 86 per infected cells with a latency of 20 minutes (Fig 1F), øEnC07 had a burst size of 40 with a latency of 20 minutes (Fig 1G), and øEnC15 had a burst size of 349 with a latency of 15 minutes (Fig 1H). Next, we verified the killing efficacy of the three phage cocktail as assessed by efficiency of plating (EOP) to evaluate how well the cocktail phages could infect other permissive hosts^31^. The results demonstrated EOPs between 0.1 and >1 for all interactions between cocktail phage and permissive hosts except for two instances: øEnA02 against CPO239 and øEnC15 against CPO053 (Fig 1I). Notably, in each case of permissive host interaction, at least one phage in the cocktail exhibited an EOP of 20% or higher.

**Figure 1.**
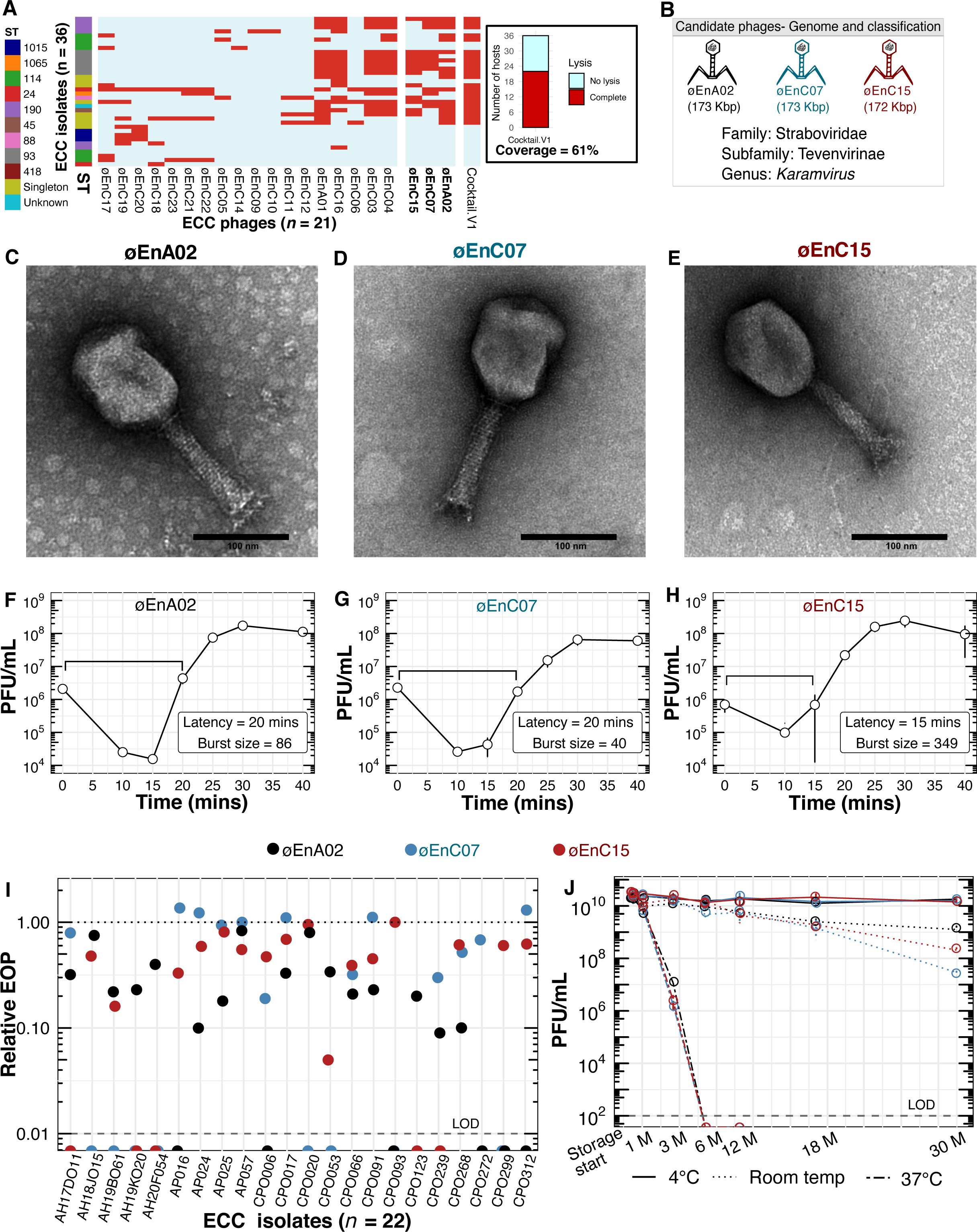
Formulation and characterisation of a three-phage combination (A-J) (A) Host range map of isolated phages (columns, *n* = 21) tested against a subset of 36 *Enterobacter cloacae* complex (ECC) isolates. Each row represents an ECC isolate, classified based on Sequence Type (ST), and is colour-coded accordingly. Singletons are STs that only occurred once. (B) Representations of phage genome size and classification of the three phages (øEnA02, øEnC07 and øEnC15) of the cocktail. (C-E) TEM images of the three phages (øEnA02, øEnC07 and øEnC15), with a 100 nm scale bar. (F-H) One-step growth curve of the three phages (øEnA02, øEnC07 and øEnC15) in the cocktail. Each point represents the average of three independent experiments, and the error bars indicate the standard error of the mean. The latency time (in minutes) and burst size (in PFU/infection) were calculated for each phage. (I) Relative efficiency of plating (EOP) values for three phages, represented by different coloured dots, across ECC isolates. The dotted line represents the EOPs on the host of isolation, which is set as 1. LOD = limit of detection. (J) Stability of the three phages (øEnA02, øEnC07 and øEnC15) in 4 °C, room temperature and 37 °C, storage conditions. LOD = limit of detection, M = months.

As we were aiming to produce a frontline antimicrobial preparation, phage stability under variable storage conditions is paramount^32,33^. We examined the stability of each phage within the cocktail under storage conditions at 4 °C, room temperature (∼22 °C), and 37 °C for 30 months and performed phage titrations at different intervals. Within the initial six months, no noticeable drop in titre occurred in lysates stored at 4 °C and room temperature, with 4 °C proving the most stable condition and maintaining phage stability for up to 30 months. Phages stored at 37 °C were the least stable, losing therapeutic titre within two months (Fig 1J). Collectively, these data suggest that these three phages have broad host range and high antimicrobial efficacy against ECC isolates, and demonstrate excellent stability, making them promising candidates for further development and evaluation.

### Identification of Phage Receptors

Bacteria can quickly evolve resistance against phage predation; this typically manifests via loss-of-function mutations in the surface-associated structures that phages adsorb to. The simultaneous use of diverse phages targeting different bacterial surface structures can minimise or delay the evolution of phage-resistance^34,35^. To identify the receptors involved in phage-host interactions, we generated phage-resistant mutants which were sequenced to identify loss-of-function mutations^36^. We identified putative phage receptors by mapping the raw reads of phage-resistant mutants to the wild-type genome. To validate these phage-receptors, we complemented the candidate wild-type (WT) gene back to its respective phage-resistant mutant and performed an adsorption assay to determine whether phage infectivity was restored. We found that each of the three phages targeted a different component of the lipopolysaccharide (LPS) structure (Fig 2A).

**Figure 2.**
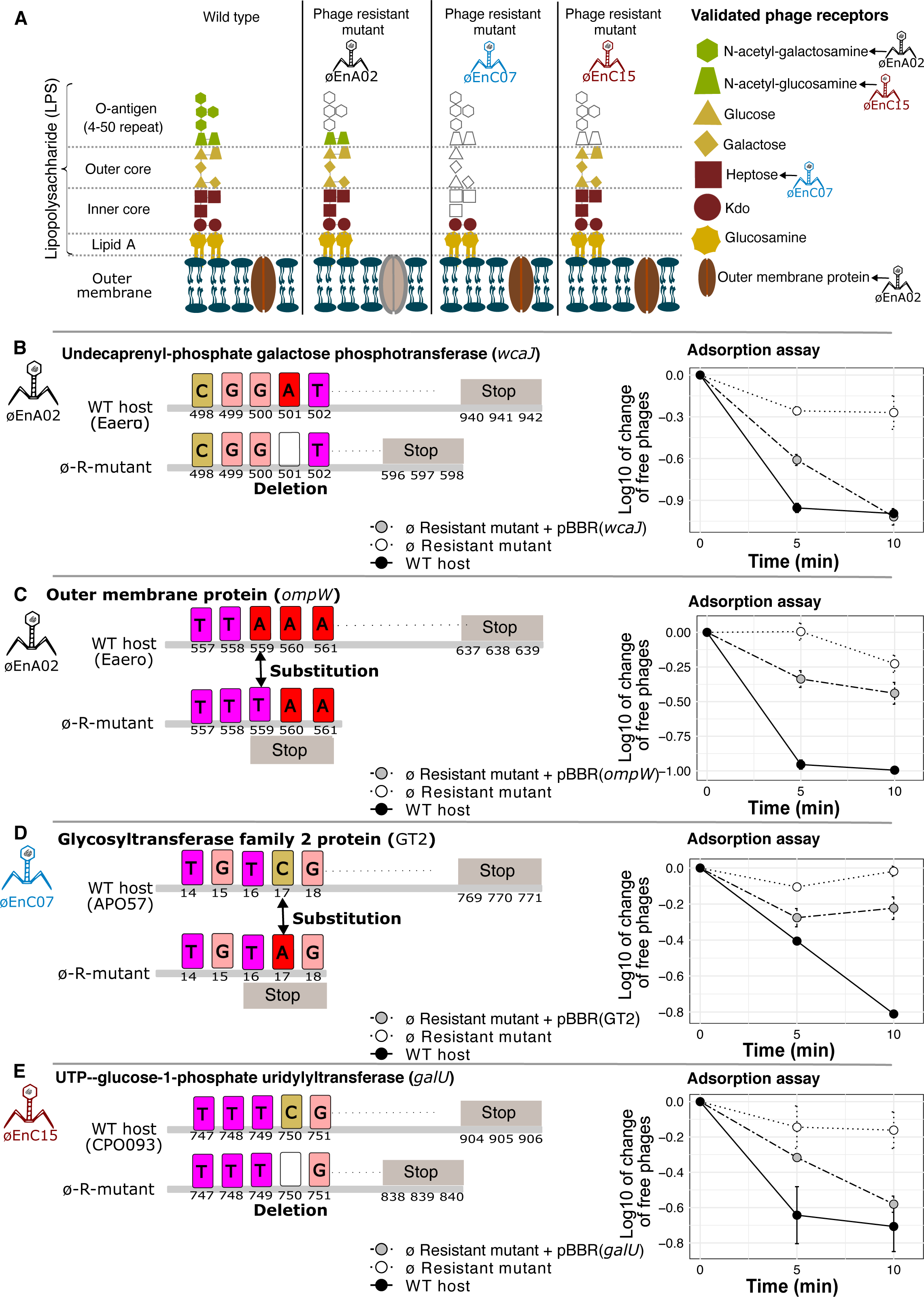
Receptors of three phages of cocktail-V1 (A-E) (A) Schematic representation of the *Enterobacter* lipopolysaccharide (LPS) and membrane structure, with validated phage receptors indicated by arrows. Colours are for illustrative purposes only and the faded section represents the potential loss of LPS structure in phage resistant mutants. Figure drawn based on^47^. (B-E) Single nucleotide polymorphisms (SNPs) identified in the genomes of phage-resistant mutants and their predicted effects. ø-R-mutant indicates phage-resistant mutant, numbers represent nucleotide positions. Graphs on the right of each panel show phage adsorption assays testing the ability of phages to adsorb to their corresponding wild-type (WT), phage-resistant mutant, and complemented hosts over 10 minutes. Each point on the graph represents the average of three independent experiments, and the error bars indicate the standard error of the mean.

In the phage resistant mutant øEnA02, we identified two loss of function mutations, the first (Fig 2B) in *wcaJ,* which is an O-antigen-associated gene^37^, and the second in an outer membrane protein (*ompW*) (Fig 2C), with both mutations resulting in the gain of an early stop codon. O-antigen is a serogroup-specific sugar-based component of the outer LPS layer of Gram-negative bacteria and is a well-characterised phage receptor^38–40^. OmpW has been shown to be a phage receptor for *Vibrio cholerae* phage^41–43^, has important roles in virulence, and acts as a colicin receptor in *E. coli*, suggesting a potential fitness trade-off associated with phage-resistance^42,43^. Following complementation, we quantified øEnA02 adsorption to the WT-host, each of the two phage-resistant mutants, and each mutant complemented with the respective WT gene. The WT-host adsorbed approximately 1 log of phage within 10 minutes, while both phage-resistant mutants showed minimal adsorption (Fig 2B and 2C). Comparatively, the complemented strains restored phage adsorption albeit only partially for the OmpW mutant (Fig 2C), suggesting that øEnA02 may use OmpW as a secondary receptor.

The øEnC07-resistant mutant had a loss-of-function mutation in a glycosyltransferase gene, which is a part of *rfa* operon and associated with LPS core synthesis^44,45^. Complementing the mutant with the WT gene restored phage infectivity and increased phage adsorption by 0.2 log (Fig 2D). Finally, øEnC15-resistant mutant had a stop codon in the UTP-glucose-1-phosphate uridyltransferase gene (*galU*), which encodes an enzyme for UDP-glucose synthesis, which is a precursor for O-antigen synthesis^46^. Complementing *galU* into the phage-resistant mutant restored phage adsorption, indicated by a 0.6 log increase (Fig 2E). The identification of phage receptors provides mechanistic insights into phage infectivity and the emergence of resistance. While all our phages broadly targeted the LPS, this was mediated through different LPS subunits, which likely reduces the emergence of phage resistance when used in combination (Supplementary Fig 3).

### In-vivo effectiveness of the preliminary phage combination (cocktail-V1)

After establishing the host range, infectivity, genomes, stability, and receptors for our preliminary three-phage combination (cocktail-V1), we evaluated its efficacy in reducing ECC burden *in vivo* using a murine model. For this model, we selected ECC isolate APO57, which was susceptible to all three phages at an EOP ∼1. We optimised the ECC inoculum dose to induce severe septicaemia in 8-weeks-old BALB/c mice, reaching an ethical endpoint within 12 hours (Supplementary Fig 4). Mice were intraperitoneally (IP) injected with 200 µL of an optimised dose of 5 × 10^6^ CFU/mL of strain APO57. One hour-post-infection (hpi), mice were treated with a single dose of cocktail-V1, which contained 10^8^ PFU/mL of each three phages at in 200 µL, while control mice received an equivalent volume of sterile PBS. The experiment concluded at 12 hpi and vital organs were collected to assess the bacterial and phage load (Fig 3A). The phage-cocktail treatment resulted in >3 log reduction (>99.9%) in the bacterial burden in the infected mouse’s blood, kidney, liver, and spleen compared to the PBS control group (*p* < 0.05) (Fig 3B). Notably, bacterial load in the blood was cleared below our detection limit (2 log) in four out of six mice. Phage load assessment revealed propagation of all three phages *in vivo*, indicating successful replication and dissemination throughout the mice (Fig 3C). We further observed that the liver and spleen exhibited higher phage concentrations compared to the blood and kidney (*p* < 0.05). However, there was no significant difference in phage concentrations when comparing the blood to the kidney or the liver to the spleen. Due to its capacity to reduce bacterial burden and ability to replicate effectively in mouse organs, we advanced the three-phage combination to the next stage of clinical evaluation.

**Figure 3.**
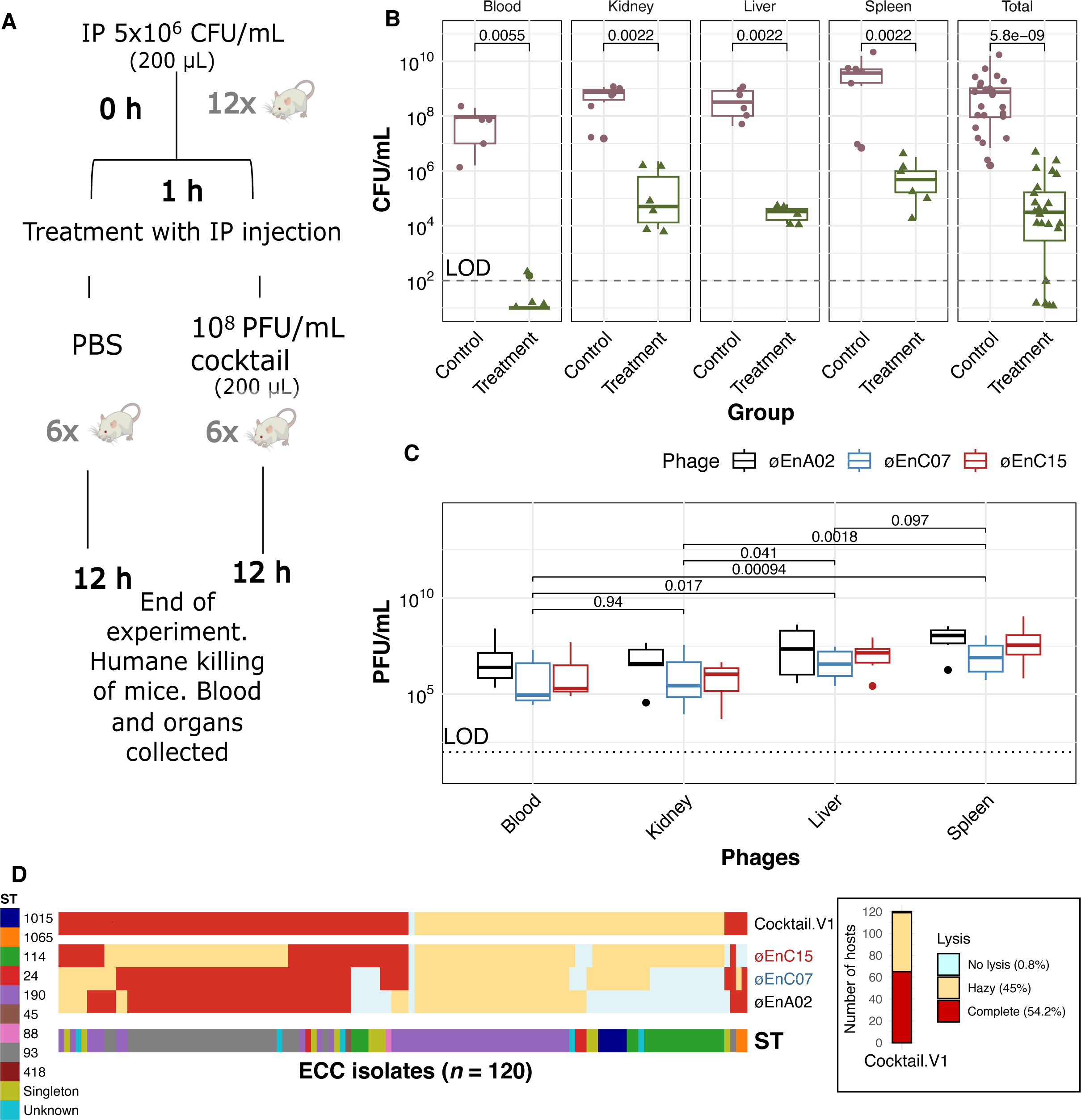
Efficacy of the phage cocktail-V1 against mouse infections and its host coverage across a range of clinical isolates (A-D). (A) Experimental scheme for the animal experiment. Grey numbers represent number of mice, black numbers represent hours post-infection. (B) The reduction of bacterial load in vital tissues of treated mice compared to the control group. Each data point represents the results from an individual mouse (*n* = 6), and the median and interquartile range are shown. Total represents data from all samples (*n* = 24) combined. Statistical analysis was performed using the Mann-Whitney U test to assess the differences between medians of bacterial count. (C) The propagation of three different phages in various tissues of treated mice. The data (*n* = 6) are shown in box plots, with medians, interquartile range, and outliers (points beyond whisker) shown. Statistical analysis was performed using the Mann-Whitney U test to assess the differences between medians of phage titre in each tissue. (D) Host range map of cocktail V1 (øEnA02, øEnC07, and øEnC15) and its individual components tested against 120 *Enterobacter cloacae* complex (ECC) isolates from The Alfred Hospital. Each column represents an ECC isolate, classified based on their Sequence Type (ST), and is color-coded accordingly. Red boxes indicate complete phage lysis of the respective ECC isolate, yellow boxes represent partial phage lysis (hazy zone), and light blue boxes indicate no observed lysis.

### Efficacy of cocktail-V1 against the wider collection of clinical ECC isolates

At this stage, we wanted to determine the effectiveness of our phage cocktail-V1 against the broader clinical collection of ECC isolates from the Alfred Hospital. We screened the lytic activity of our phage cocktail and its individual components via a spot assay against 120 clinical ECC isolates (Fig 3D). At the time, this represented the entire ECC collection at The Alfred, encompassing a diverse range of sequence types (STs) that contributed to the nosocomial outbreak. Two researchers who were blinded to the experiment visually scored the zones on the bacterial lawn as complete lysis (phages clearing the bacteria), hazy (bacteria partially cleared) or no lysis (no effect of phages) (Supplementary Fig 5). The three-phage cocktail completely lysed 54.2% of the 120 ECC isolates, which was a decrease of ∼7% in lysis when compared against the initial 36 isolate collection cocktail-V1 was built upon (Fig 1A). Hazy spots (∼45%) were not confirmed for productive infection via the soft overlay (top) agar assay, but likely represented a combination of low EOP, phage resistance and bactericidal activity resulting from “killing from without”^48^. We also observed association between phage susceptibility and ST of the ECC isolates, with ST-114, 190 and 1015 being the least susceptible to the cocktail. The modest host range coverage of 54.2%, along with limited activity against specific STs, underscored the need for a more tailored approach to phage isolation and the refinement and expansion of phage infectivity to ensure comprehensive coverage across broader STs encompassing the ECC nosocomial outbreak.

### Phage training to improve adaptability and infectivity

A fundamental difference between phages and antibiotics is phages can evolve and adapt to changes in their hosts. This can be harnessed to train phages and achieve improved fitness and therapeutic efficacy^49–51^. Using an experimental evolution approach, we set out to improve the lytic capacity of the three phages from cocktail-V1. Our first goal was to determine the optimum phage training duration that would result in improved lytic capacity. We selected ECC strain AH17D011 and øEnC07, a phage-host pair with a moderate EOP of 0.79. Following 24 hours of propagation, we purified the population of trained phages, which were then used to infect a naive host population, with the entire process being repeated for 10 days. After five days, we performed a growth kinetics assay to determine the lytic efficacy of the phages and calculated phage scores as an indicator of phage fitness^52,53^. We observed that the phage score increased sharply starting on day five of training and plateaued afterwards (Supplementary Fig 6). Based on these observations, we determined that the optimum phage training period was between five to seven days, with seven days being selected for further experiments. Importantly, following the seven-day phage training, evolved phages were twice single-plaque purified to ensure that a single phage genotype was taken for downstream characterisation, including lytic activity via EOPs, growth curve, and genomic changes (Fig 4A).

**Figure 4.**
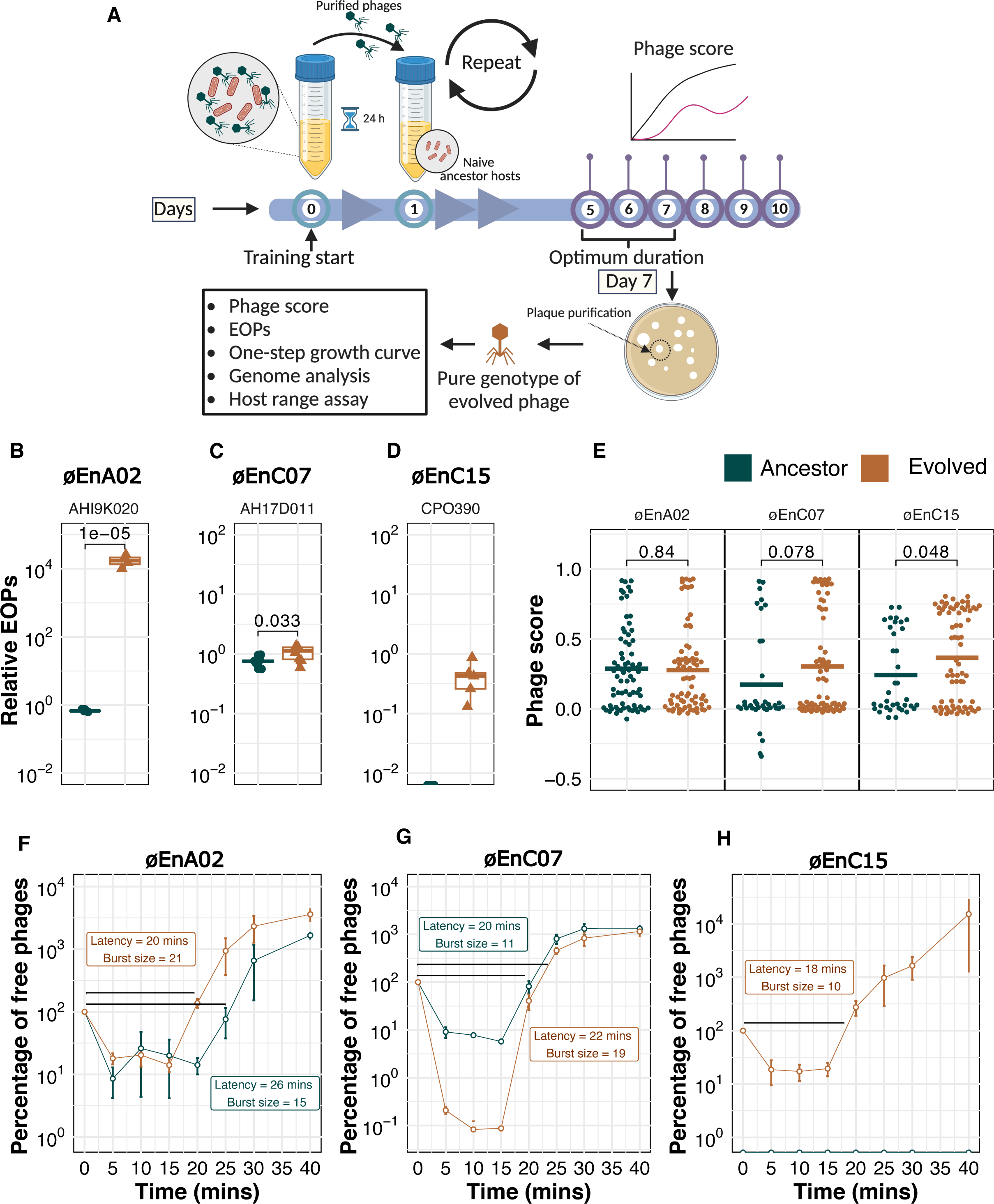
Phage training and its effect on phage adaptation (A-H). (A) Experimental protocol for phage training. Ancestral phage was propagated with training hosts for 24 hours. Subsequently, the purified phage population was repropagated on a naive population of the same training host, with this cycle repeated for 10 days. After five days, daily measurements of phage growth curves were conducted using the evolved population to find the optimum training duration. The final evolved phages (i.e., day 7) were isolated via double single-plaque purification to obtain an individual genotype of phage mutants. Then, the growth characteristics of these single genotype evolved phages were assessed. (B-D) The relative efficiency of plating (EOP) of evolved phages (øEnA02, øE2nC07, and øEnC15) compared to their ancestral counterparts. The labels on top of the boxes represent the training ECC host. Each point (*n* = 6) represents data from independent biological replicates. *p* values were calculated by a two-tailed t-test with Welch’s correction. Box bounds indicate 25th and 75th percentile. (E) Phage score of ancestor phages compared with their respective evolved phages, tested against a subset of 36 *Enterobacter cloacae* complex (ECC) isolates. Each dot represents a phage score (*n* = 36). *p* values were calculated by a two-tailed t-test with Welch’s correction. (F-H) One-step growth curve of evolved phages (øEnA02, øE2nC07, and øEnC15) compared to their ancestral counterparts, on their respective training hosts. Each point represents data from independent biological replicates (*n* = 3), and the error bars indicate the standard error of the mean.

We then examined whether phage training could enhance lytic activity against hosts that had lower EOPs and were less responsive to a specific phage. As such, we trained the remaining two phages from our cocktail: øEnA02 with host AH19K020, (EOP 0.2), and øEnC15 with the apparently non-permissive host CPO390 (EOP below the limit of detection). Following phage training, EOPs improved by 4,000-fold for øEnA02 (*p* <0.001) and 8.7-fold of øEnC07 (*p* = 0.033) against their training hosts. Intriguingly, øEnC15, initially incapable of productive infection on its training host, was found to infect it and propagate therein at an EOP of 0.4 after training (Fig 4 B-D). To evaluate the broader fitness of trained phages and any potential trade-offs, we compared their lytic activity to that of their ancestral counterparts across our 36 ECC isolate subset. The evolved phages showed an increased average phage score for øEnC07 (*p* = 0.078) and øEnC15 (*p* = 0.048), while øEnA02 showed minimal change (*p* = 0.84) (Fig 4E, Supplementary Fig 7).

To determine whether the observed improvements in the killing efficiency of the evolved phages were accompanied by changes in their life cycles, we conducted one-step kill curves on their training hosts. Compared to its ancestor, the evolved øEnA02 had a faster replicative cycle as demonstrated by a shorter latency period (20 min vs 26 min) and released more progeny virions (burst size; 21 vs 15) (Fig 4F). Evolved øEnC07 showed an enhanced adsorption rate, as indicated by the steeper decline of free phage percentage in the first 15 minutes of infection, with a comparable latency (22 min vs 20 min) and larger burst size (19 vs 11) (Fig 4G). As for the øEnC15, we lacked an ancestral phage that could infect the host for the comparison, so we evaluated the life cycle parameters of the evolved generation, revealing a latent period of about 18 min and a burst size of 10 (Fig 4H).

To explore the molecular mechanisms underlying the improved phage efficacy, we conducted genome sequencing and comparative analysis between the evolved phages and their ancestors (Supplementary Fig 8). We identified several SNPs in the evolved phages, with the majority located in the tail region of the genome, including the tail fibre and receptor-recognising proteins. These proteins are critical for the initial adsorption of phages to host bacteria. Interestingly, we found that some of the SNPs were unique to certain phage-host pairs, indicating the specificity of the evolutionary process. Based on our observation, we hypothesised that the improved phage efficacy resulted from selection and accumulation of beneficial mutations during the training process, which enabled the phages to better recognise and bind to their hosts. Collectively, these results demonstrate that our phage training approach successfully improved phage infectivity against inefficient or even non-permissive hosts, via mutations in phage tail structures, impacting divergent variables of the phage lifecycle and enhancing killing efficiency.

### Targeted phage isolation against problematic sequence types (STs)

Thus far, we focused on characterising three phages of our cocktail-V1 and demonstrated their *in vitro* and *in vivo* efficacy, followed by phage training to improve their lytic capacity on low-EOP hosts. However, this three-phage combination had limited effectiveness against specific STs from the ECC collection (Fig 3D). To address this, we selected ECC strains from STs that lacked sufficient phage coverage as hosts for targeted phage isolation. Five novel phages were isolated, and their activity tested via spot assay (Supplementary Fig 9). Based on the host range map and complementary spectrum of activity provided by cocktail-V1, we selected two candidate phages (øNando and øTaquito) for further characterisation (Fig 5A). Genomic and taxonomic classification revealed both phages belonged to the Genus *Pseudotevenvirus* under the Straboviridae family (Fig 5B). TEM confirmed the presence of an icosahedral head (70-75 nm diameter) and a long contractile tail (210 nm), consistent with the characteristics of the Straboviridae family (Fig 5C and 5D). The genomes of both phages lack transposase or integrase genes, antimicrobial resistance markers, and virulence genes, indicating that they are not temperate and are suitable candidates for therapeutic use.

**Figure 5.**
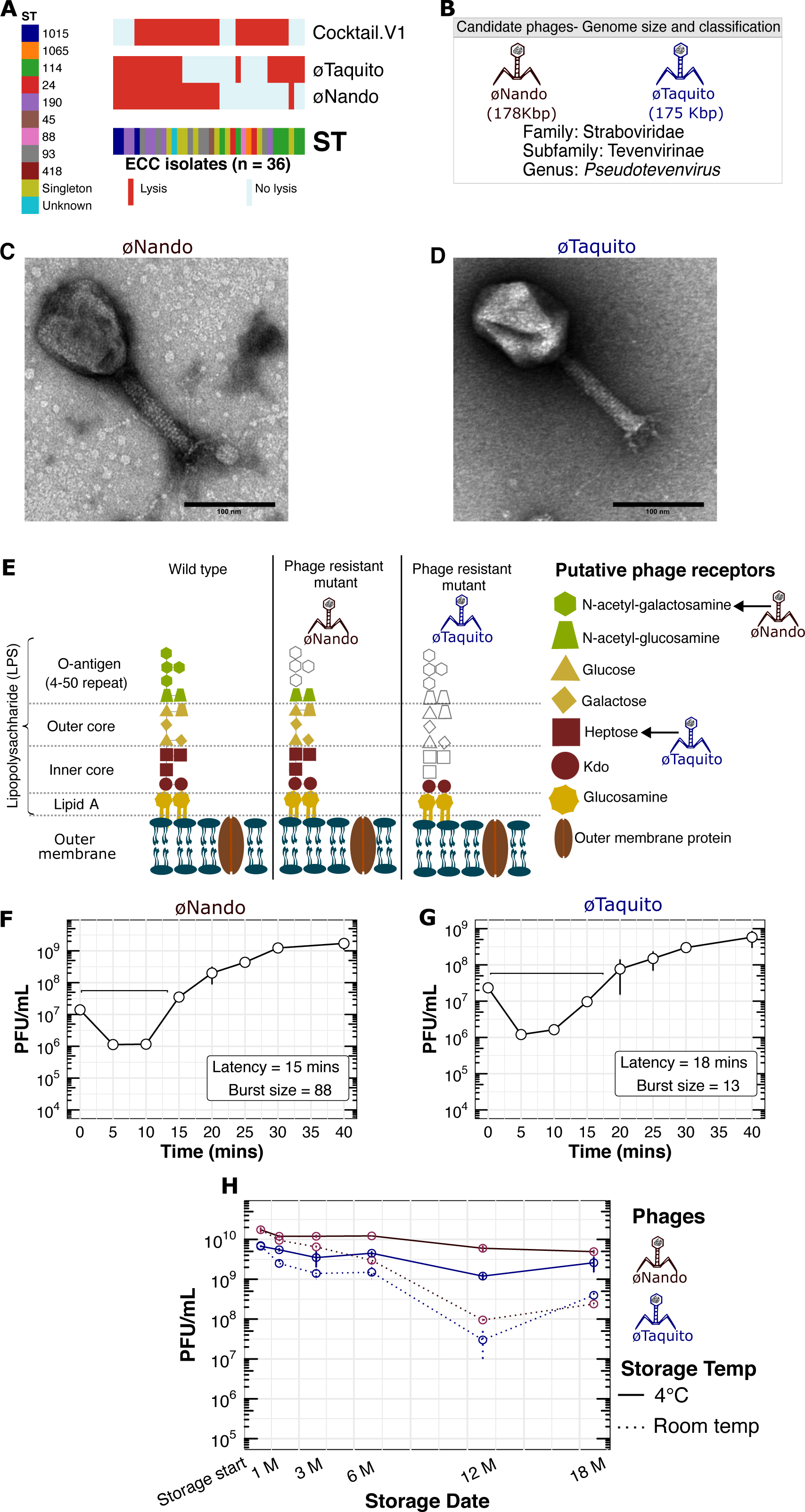
Targeted phage isolation and expansion of host range (A-H). (A) Host range map of isolated phages and cocktail-V1 tested against the initial subset of 36 *Enterobacter cloacae* complex (ECC) isolates. Each row represents a phage. Sequence Types (ST) are colour coded. Singletons are STs that occurred once. Red boxes show phage lysis and light blue boxes show no lysis. (B) Representations of phage genome size and classification of the phages (øNando and øTaquito). (C and D) TEM images of the two phages. Scale bar = 100 nm. (E) Schematic representation of the *Enterobacter* lipopolysaccharide (LPS) and membrane structure, with putative phage receptors indicated by arrows. Colours are for illustrative purposes only and the faded section represents the potential loss of LPS structure in phage resistant mutants. Figure drawn based on^47^. (F and G) One-step growth curves of øNando and øTaquito. Each point represents the average of three independent experiments, and the error bars indicate the standard error of the mean. (H) Stability tracking of øNando and øTaquito over 18 months, with storage at 4 °C (solid line) and room temperature (dotted line).

Identification of receptors revealed that both phages targeted different bacterial cell surface structures to mediate infection (Fig 5E). For the øNando-resistant host (mutated from strain CPO448), we found a loss of function mutation within *wzzB*, which encodes a protein involved in the biosynthesis of O-antigen and helps determine the chain length^54^. Additionally, for øTaquito, we discovered a loss-of-function mutation in a resistant variant of host CPO165 within the *rfaQ* gene. RfaQ is involved in LPS biosynthesis as it transfers the first two heptose residues in the inner core of LPS, and its loss leads to a severely truncated LPS alongside pleiotropic effects on bacterial cells^44^. We next evaluated one-step growth curves for both phages, with øNando demonstrating a burst size of 88 and a latency of 15 minutes (Fig 5F) and øTaquito a burst size of 13 and a latency of 20 minutes (Fig 5G). Regarding stability, both phages when stored at 4 °C maintained stability for up to 18 months (Fig 5H), whilst room temperature storage supported phage stability for at least six months. These findings, combined with our initial phage results, warranted the inclusion of these two additional phages to our cocktail aiming towards an expanded host range with coverage against problematic STs that cocktail V1 did not target.

### Improved five-phage cocktail achieved broad coverage against clinical ECC isolates

Our first-generation, three-phage cocktail (cocktail-V1) achieved a modest ∼54.2% host range coverage against the ECC isolates endemic to The Alfred Hospital. To improve its coverage and efficacy, we trained the original three phages to enhance their lytic activity against selected low-EOP hosts, followed by a targeted isolation of two novel phages against problematic STs. Our focus then shifted to identifying the optimal combination of these phages for the development of an improved cocktail. To this end, we prepared eight combinations of phages comprising three-, four-, or five-phage combinations from the above mentioned eight candidate phages (three ancestral phages, three evolved phages, and two targeted-isolation phages). We tested these cocktails and PBS control (nine samples labelled A-I) using a spot assay against an expanded collection of 156 clinical ECC strains, which included our previous panel (*n* = 120) (Fig 4D) plus 36 additional strains that caused infections in the hospital during the timeline of this study.

Both of our five-phage combinations: combination G utilising evolved phages (ev_øEnA02, ev_øEnC07, ev_øEnC15, plus phages øNando and øTaquito) and combination I with ancestral phages (øEnA02, øEnC07, øEnC15, øNando and øTaquito) produced complete lysis on spot assays against ∼65% and 75% of the collection, respectively. These ranges included isolates from ST 114 and other previously untargeted STs, therefore surpassing the performance of other combinations (Fig 6A; Supplementary Fig 10). This marked a substantial improvement over cocktail-V1’s 43.9% spot lysis on the expanded collection of strains. Next, to resolve whether hazy spots were indicative of productive infection, we used growth kinetics assays and calculated phage scores. We selected 67 hosts for analysis, including all hosts where either both or any of combinations G and I created hazy spots, and a few hosts with complete lysis as controls. To distinguish productive infections, optimum cut-off value of 0.28 in phage score was determined using the maximum likelihood estimation^55^. The majority of the hosts with hazy zones on spot assays yielded productive infections except for 12 instances in cocktail G and 9 in cocktail I. All instances of complete lysis spots were confirmed to be productive, except for one in combination G. Additionally, no difference in the phage score was observed between G and I (Fig 6B). Combining these results with the spot assay data, we deduced that both G and I achieved a lysis rate of at least 92% (144 out of 156 isolates) (Fig 6C). However, given our previous demonstration of enhanced fitness of the evolved phages compared to their ancestral counterparts (Fig 4E), combination G was selected for manufacturing of stable, therapeutic-grade phage product.

**Figure 6.**
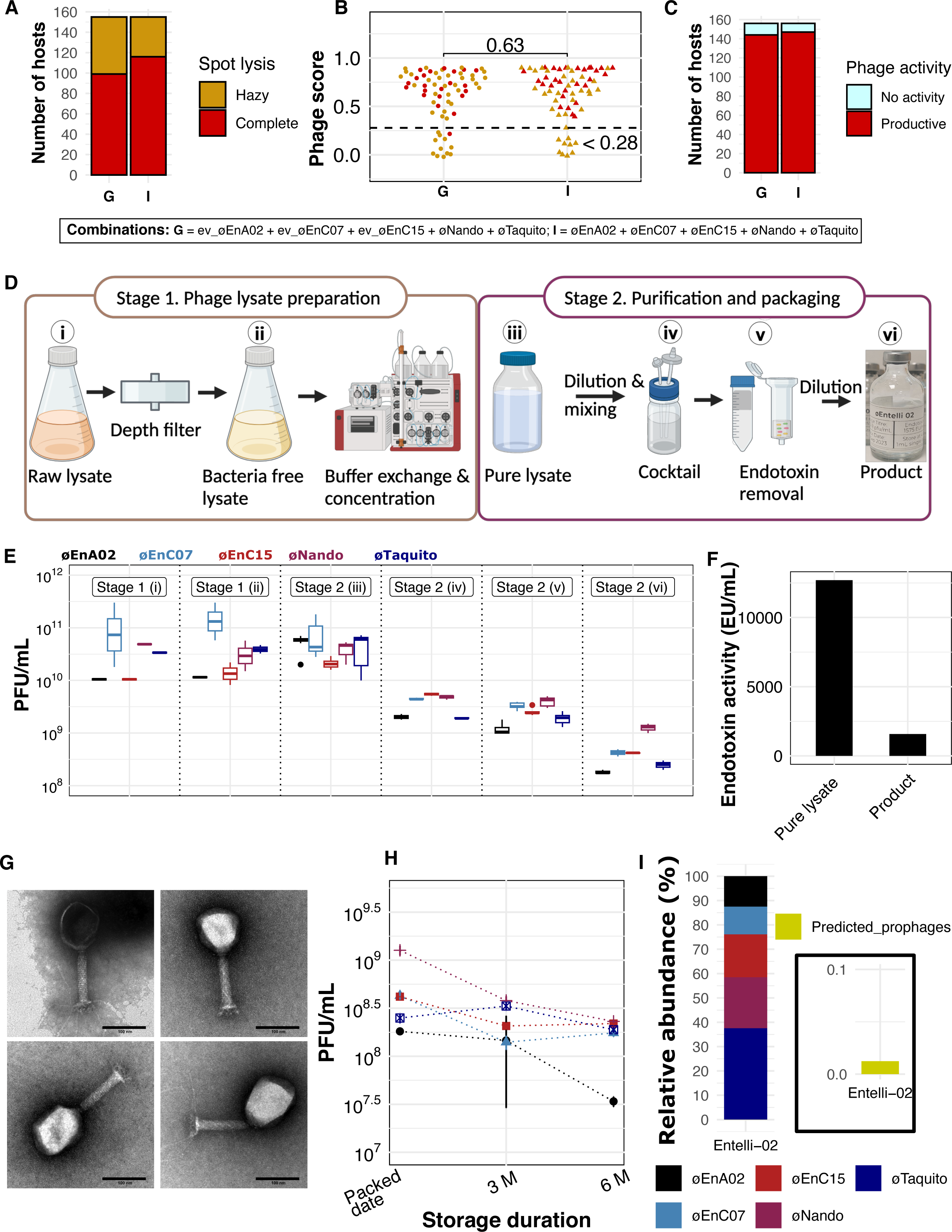
Phage combination screening and manufacturing and quality control of phage product (A-I) (A) Host range coverage of phage combinations (G and I) tested against 156 *Enterobacter cloacae* complex (ECC) isolates based on spot assay data. (B) Comparison of phage score of combinations G and I against *Enterobacter cloacae* complex (ECC) isolates. Each dot represents a phage score (*n* = 67). *p* values were calculated by a two-tailed t-test with Welch’s correction. Dotted line (<0.28) indicates the threshold for characterising a phage activity as productive. (C) Host range coverage (*n* = 156) of phage combinations (G and I), based on combined phage score and spot assay data. (D) Schematic representation of the therapeutic-grade phage production protocol. (E) Phage titre at each production steps stage. Each data point represents results from a replicate (*n* ≥ 3), whiskers indicate maximum and minimum values, box bounds indicate 25th and 75th percentile, with centre line indicating the median. (F) Endotoxin activity as Endotoxin Unit (EU) per mL of pure lysate [stage 2(iii)] and the final product [stage 2(vi)]. (G) TEM images of phage product (*Entelli-02*) showing intact virions and no contaminants. (H) Phage titre stability of the packed product on storage at 4 °C over six months. (I) Normalized relative coverage (relative abundance) of sequencing reads from the whole genome of *Entelli-02* to component phages and predicted prophages.

### Production of a therapeutic-grade phage cocktail *Entelli-02*

We manufactured a therapeutic-grade phage product that we named “*Entelli-02*” from the five-phage combination (G) at our in-house facility, the Monash Phage Foundry (Fig 6D). In a first stage, we produced each phage individually. Here, phages were amplified through overnight propagation with their respective ECC hosts of isolation. The resulting phage lysates were processed using a two-stage sequential depth filtration (0.5-15 µm and 0.2-3.5 µm retention ratings) to reduce bacterial biomass followed by sterilising grade (0.2 µm) filtration into sealed glass containers. Bacteria-free lysates (approximately 1 L each) were then transferred to a clean-room facility for further processing. Phage lysates were diluted ∼10-fold in 1X PBS followed by buffer exchange and concentration using tangential flow filtration (TFF), resulting in a recovery between 130 -160 mL of washed and concentrated lysate from each production run. We observed a recovery efficiency ranging from 24% to 76%, with a PFU/mL count exceeding 10^10^ PFU/mL for each phage (Fig 6D) (Supplementary Table 1).

In stage two of production, we mixed and, if necessary, diluted the five concentrated lysates to achieve a uniform phage product with a titre of >10^9^ PFU/mL per phage in a total of 75 mL. We then depleted endotoxin using both EndotrapHD and 1-Octanol treatments, resulting in nearly 10-fold reduction in endotoxin levels (Fig 6E). Finally, we diluted the cocktail ten-fold in 1X PBS to obtain our final product consisting of an average titre of ∼5 × 10^8^ PFU/mL/phage in a total volume of 500 mL. The cocktail was twice filter-sterilised using sterilising-grade filters (0.2 µm), followed by quality control validation for phage titre and endotoxin levels (see detailed methods). The cocktail was then packaged into syringe-accessible vials, each containing 35 mL of *Entelli-02*, which is suitable for a two-week treatment course with an effective dose of 1 mL administered twice daily (b.i.d.). The final packed product underwent external validation for sterility and endotoxin, according to the USP71 and USP85 guidelines, respectively, with >10% of the production batch sent for validation. The final product contained no visible growth of microorganisms and had an endotoxin concentration of 1,575 EU/mL, which according to FDA guidelines (5 EU × kg × h) would be safe for intravenous administration b.i.d. to a patient >30 kg^56^. Electron microscopy images confirmed the visual integrity and cleanliness on *Entelli-02* (Fig 6F). Furthermore, considering these phages were amplified on clinical isolates, we also performed whole genome sequencing of the final product, followed by read mapping to the individual phage genomes, the host bacterial genomes, and predicted prophage regions (see Methods). Sequence analysis showed that >99.2% of reads mapped to the phage genomes, 0.27% to the bacterial genome, and about 0.08% to predicted prophage regions (Supplementary Table 2). To further, examine the prophage presence, we determined the normalised coverage per million reads (relative abundance) of five phages and predicted prophages. Considering the average titre of ∼5 × 10^8^ PFU/mL/phage in our product and based on the relative abundance of 0.7 (0.01%) of prophage genomes (Fig 6), we inferred that the level of prophage in *Entelli-02* is <5 × 10^4^ PFU/mL. Finally, an important aspect of the chemistry, manufacturing, and control process is maintaining the stability of the individual components over time. We measured titre of the individual phages using selective platting within our final *Entelli-02* product and found no indication of stability issues over 6 months of storage at 4 °C (Fig 6G). In summary, these tests confirmed the suitability of *Entelli-02* for compassionate-use and rapid frontline deployment for the treatment of ECC infections originating from The Alfred Hospital.

## Discussion

Currently, the lack of standardised and readily available phage products for use against nosocomial multidrug-resistant pathogens is a major impediment to administering effective phage therapy in healthcare settings^57^. Here, we employed a unique approach that leverages the utility of broad-spectrum phage cocktails with the increased efficacy and specificity found in personalised phage therapy. We define this approach as an ‘institutional cocktail’ that has been designed upon a high-quality and representative collection of nosocomial pathogens, and that can be employed locally and rapidly as a frontline therapeutic with high probability of antimicrobial activity. Our integrated academic-clinical approach was personalised towards the endemic pathogen profile at the institutional level and aimed to improve both the treatment efficacy and response time for future compassionate-use phage therapy cases. Through this, we developed an institutional cocktail, which we named *Entelli-02,* to serve as a frontline therapeutic against AMR *Enterobacter cloacae* complex (ECC) infections at The Alfred Hospital, in Melbourne, Australia. We envision this approach being employed to treat other nosocomial pathogens, expanded across additional institutions, and iterated upon to improve product efficacy towards an everchanging pathogen population.

Phage therapy has made significant progress in recent years. Personalised phage therapy approaches have reported clinical improvement in ∼75% of patients, bacterial eradication in ∼60-80% of patients, and adverse events in ∼15% of patients^8,15^. However, personalised phage therapy applications are time-consuming, complex to administer, and difficult to scale, which limits the number of patients that can be treated. Conversely, broad-spectrum phage cocktails designed to treat entire pathogen complexes as stand-alone therapies have been produced under current pharmaceutical GMP-guidelines and can be made rapidly available. Yet the performance of these defined cocktails in randomised controlled trials completed to date has been underwhelming^2,8–12^, hinting at a disconnect between the strains used to develop these products and the clinical strains in patient cohorts being treated. While the mixing of phages that target multiple strains of a single species or even multiple species is a traditional and widely used method, there are several factors that can compromise the efficacy of such broad-spectrum cocktails^8^. For instance, building the cocktail using limited culture collections or the use of clinically irrelevant hosts may result in poor coverage and inefficient killing. Additionally, the laboratory-based phage-host match may not necessarily correlate with the lytic activity of the phage *in vivo*, and bacteria can quickly evolve phage resistance. Importantly, few broad-spectrum cocktails identify the phage-receptors their constituents use^58^, with the simultaneous use of phages targeting the same receptor possibly resulting in phage competition for adsorption, further reducing the cocktail’s efficacy. Thus, it is important to implement a rational approach for the selection of phages in a cocktail, ensuring a broad host range, high efficacy, and minimal risk of resistance development^58–61^.

For the iterative development of *Entelli-02*, we began by isolating phages against a sub-collection of 36 genetically diverse isolates of ECC, representative of the broader endemic ECC outbreak at The Alfred Hospital. From this initial phage library, we formulated a first-generation three-phage combination (cocktail-V1) based on orthogonal host range coverage and lytic activity. We then comprehensively characterised each phage for therapeutic relevance, which included conducting one-step growth curves, efficiency of plating (EOP), stability, genome sequencing, phage receptor validation, and *in vivo* antimicrobial efficacy. The observed 3 log_10_ reduction of ECC in our *in vivo* mouse model indicates high efficacy of cocktail-V1. Furthermore, these phages appear to use at least two different receptors, OmpW and LPS, and importantly, different subunits of the LPS molecule. This target receptor diversity makes the combined use of these phages less likely to result in inter-phage competition and selection for phage-resistance. This latter point was evidenced by the apparent propagation of all three phages in our *in vivo* mouse model (Fig 3C). Screening this first-generation cocktail against the wider clinical collection of 120 ECC isolates revealed a moderate host range coverage of 54%, with low killing efficiency against a few particular STs. Based upon these results, we decided to improve our phage cocktail by iterating upon our experimental design through two different approaches: first, a phage training protocol to improve the lytic activity of our original phages and, second, the targeted isolation of novel phages against those low-coverage STs.

To improve the killing efficiency, we employed an experimental evolution approach, more commonly known as phage training ^49–51^. Phage training would help improve the adaptability of phages to other hosts, while hopefully maintaining their capacity to infect original hosts ^62–65^. Post-training, we isolated and characterised single phage genotypes from the evolved populations. We emphasise the importance of isolating and characterising well defined, singular phage genotypes from phage training techniques. It’s essential to only use known phage genotypes in therapy to ensure it’s safe and meets regulatory requirements. Selection of trained phages was based on improved phage score^66^, which eventually showed improvement of EOPs, improved adsorption, latency, and burst size, compared to untrained ancestors. Notably, even in a case of a seemingly non-permissive host, a trained phage adapted to it, resulting in efficient lysis. Coevolutionary interactions between bacteria and phage usually involve adaptations to the initial adsorption of the bacteriophage to its hosts, often through changes in its tail structure ^67^. As expected, we observed mutations within phage tail fibre genes in our trained phages. Notably, we also identified mutations in other structural genes, which could impact the phage lifecycle and warrant further investigation. By combining phage training with careful phenotypic selection, we improved the killing efficiency of our phages. To broaden the spectrum of cocktail-V1, we employed targeted phage isolation against clinical ECC strains with STs not covered by the initial cocktail. This expanded our phage library and allowed us to select two additional phages that targeted different regions or potentially different variant of the O antigen in LPS^68^. This selection complemented the host range of cocktail-V1. Similar to the cocktail-V1 components, these phages exhibited high antimicrobial efficacy *in vitro* and lacked lysogeny and virulence-related genes. Their inclusion enhanced our cocktail’s ability to target elusive STs, resulting in a widened host range practically covering the entire ECC collection from the Alfred, demonstrating the importance of optimising and personalising phage cocktails for specific pathogens and epidemiological niches. We also emphasised the importance of regularly updating the phage cocktails based on the evolving genetic background of clinical isolates to ensure the best efficacy in phage therapy.

*Entelli-02*, our second version phage cocktail, was selected by screening different candidate combinations of phages against the entire cohort of ECC isolates via spot and bacterial growth kinetics assays. The best performing combination comprised our three evolved phages plus the two phages isolated against targeted STs (Fig 6A). While the host range of this combination was marginally narrower than the combination containing the three ancestral phages, the evolved phages showed higher adaptability to hosts, as evidenced by decreased latency period, increased burst size and adsorption, which resulted in higher phage scores for two out of three evolved phages (Fig 4E). Furthermore, evolved phages, trained on selective hosts, maintained their infectivity across the broader collection of ECC isolates, indicating no fitness trade-off for the evolved phages.

In preparation for clinical administration, we manufactured a clinical-grade *Entelli-02* and ensured a final stable product. Our production pipeline maintained high recovery efficiency and implemented traceable manufacturing procedures, ensuring consistency and reliability throughout the manufacturing process. The conclusive assessment of *Entelli-02*’s wide host coverage (∼92%) on clinically relevant ECC strains further substantiates its clinical potential and capacity for rapid deployment. In summary, our study represents a comprehensive and multifaceted approach to generating an ‘institutional cocktail’ *Entelli-02* that bridges the gap between personalised phage therapy and broad-spectrum phage cocktails. Developed in response to a local hospital’s outbreak and based on a clinically curated strain collection, *Entelli-02* is designed to combat an endemic AMR pathogen in a nosocomial setting as a frontline therapeutic. Importantly, this ‘institutional cocktail’ offers a streamlined alternative to the extensive processes typical of personalised phage therapies, which leverages advantages from both single-patient compassionate use and broad-range phage cocktails, setting the blueprint for establishing ‘institutional cocktails’ against other AMR pathogens across different hospitals and institutions. Further work is needed to determine the breadth and activity of our ECC phage cocktail against both national and international ECC isolates, including conducting clinical trials to evaluate its efficacy and safety in treating AMR ECC infections at The Alfred Hospital. Finally, this approach can be iterated and adapted to the changing genetic background of ECC clinical isolates at The Alfred Hospital, allowing us to produce future improved versions of the *Entelli-02* cocktail.

## Methods

### *Enterobacter cloacae* complex (ECC) clinical isolates

This study used clinical isolates of *Enterobacter cloacae* complex (ECC) from The Alfred Hospital, Melbourne, Australia, collected between 2010 and 2021. In total, the repository at The Alfred Hospital contained 206 bacterial strains. For initial phage isolation and analysis, we selected a subset of 36 strains to represent the sequence type (ST) diversity of whole collection set. To further the scope of our investigation, the study was expanded to include up to 156 isolates to evaluate phage and cocktail activities. No patient data were collected or used in these experiments and all strains were retrieved without identifiable patient data. Strain details are listed as Supplementary Data 1.

### Bacterial strains, plasmids and growth conditions

We used lysogeny broth (LB) - 10 g/L tryptone (Oxoid Ltd, Basingstoke, UK), 5 g/L yeast extract (Merck Life Science Pty Ltd; Darmstadt, Germany), 10 g/L NaCl (Merck) to grow bacteria, unless otherwise mentioned. Agar (Merck) at a concentration of 15 g/L was employed for agar plates and 7.5 g/L for the creation of a soft agar double-layer (top agar). All bacterial cultures were grown aerobically at 37 °C overnight. We used Omnipur phosphate-buffered saline (PBS) (Merck) for all experiments unless otherwise mentioned. In gene transformation experiments, we used vector pBBRMCS^69^ and *E. coli* DH5-α competent cells (C2987H, NEB—New England BioLabs Inc., Ipswich, USA) as a host.

### Phage isolation and purification

Phages were isolated from environmental sources through enrichment process^70^. Briefly, 3-4 batches of raw sewage were combined in a flask, totalling up to 100 mL, followed by addition of 10 mL of 10× LB along with supplements of 1 mM CaCl_2_ and MgCl_2_. Next, 1 mL of overnight cultures of up to three targeted bacterial hosts strains were added into the mixture and incubated at 37 °C overnight. The resulting lysates were purified through centrifugation, filtration, and chloroform treatment^71^. Phage activity was accessed by spotting 10 µL of lysates onto bacterial lawns on soft agar. When bacterial lysis was observed, we plated serial dilutions of the lysate with the host bacteria using the soft overlay (top) agar method and examined for clear plaques on the bacterial lawn after overnight incubation. For two successive rounds, plaques were individually picked from each plate, resuspended in 500 μL of PBS by vigorous vortexing, and reamplified. Lysates prepared from the purified plaques were quantified in plaque-forming units (PFU) per mL and stored at 4 °C.

### Determination of host range, efficiency of plating, one-step growth curve and adsorption assay

Host range was examined by spot assays. We applied 10 µL of phage lysate (≥10^8^ PFU/mL) onto a bacterial lawn prepared by mixing 1 mL of bacterial culture with 3 mL of soft agar, and spreading over an LB agar plate. For the initial host range assessment on the host subset (*n* = 36), spot assay results were further confirmed for their ability to form plaques, verifying productive infection. For screening in broader sets (*n* = 120 and *n* = 156), the resulting zones were recorded as clear (phage clearing bacteria), hazy (bacteria partially lysed), or no lysis (no activity). Three researchers assessed these results blindly to each other and the phage combinations used (Supplementary Fig 5). Efficiency of plating (EOP) was examined by comparing PFU/mL on the host of interest to PFU/mL on the original isolation host. Phage lysates were serially diluted, mixed with each host and plated using top agar to determine the PFU/mL. For one-step growth curves, bacteria from overnight broth cultures were diluted 1:50 in LB and grown until an optical density (OD_600_) of 0.2 was reached (corresponds to ∼10^8^ CFU/mL) (approximately 2 h). The culture was then infected with phage at a multiplicity of infection (MOI) of 0.1. Phage was allowed to adsorb for 5 min at 37°C with orbital agitation at 190 rpm. The mixture was then pelleted (4,000 × g, 2 min, room temperature), resuspended in fresh, pre-warmed LB broth, and incubated. Samples (100 µL) were collected every 5 or 10 mins for 1 h, serially diluted in PBS, and plated for phage quantification. The experiment was repeated in at least three different occasions. Using these data, we calculated the latent period and burst size as described previously^72^. Adsorption assays were performed based on the principles of a one-step growth curve, with some modifications. Phages were mixed with their bacterial hosts at an MOI of 0.1 and samples were taken at 0, 5 and 10 mins. Each sample was immediately diluted 1:10 in PBS and centrifuged to remove adsorbed phages; the supernatant was plated for phage quantification, and calculation of the changes in free phage concentration.

### Isolation of phage-resistant mutants

To generate phage-resistant mutants, we followed the broth culture method^35^. First, we incubated phages with their respective isolation host at an MOI of 1 in a 96-well microplate, with a total volume of 200 μL each, in nine replicates. After overnight incubation, we transferred 20 μL of the resulting growth to fresh LB containing 10^6^ PFU/mL of phage particles. We repeated this process for three consecutive days.

After three days, we purified the resistant bacterial colonies through two rounds of single-colony isolation and confirmed phage-resistance by top agar assays using the phage-resistant mutants and their respective phages, expecting an absence of plaques after incubation. Subsequently, we performed adsorption assays to further confirm phage resistance. Stocks of the verified mutants were stored at –80°C until further experimentation.

### Transmission electron microscopy (TEM)

To examine through TEM using negative staining, phage lysates were purified using Vivaspin 6 centrifugal concentrators (MWCO 1,000,000 kDa; Merck). The lysate was washed several times with SM buffer (100 mM NaCl, 8 mM MgSO4·7H2O, 50 mM Tris-Cl in water). The cleaned lysate was then applied onto a copper TEM grid (200 mesh, SPI, USA) with a carbon-coated ultrathin formvar film. A 10 µL droplet of phage suspension was placed on the grid, left undisturbed for 30 seconds, and then dried using filter paper. Next, a 10 µl droplet of uranyl acetate water solution (1% w/v) was added to the grid surface and left for 20 seconds. After drying the grid with filter paper, it was examined under TEM (JEM-1400 Plus; Jeol, Japan) operating at an accelerating voltage of 80 kV.

### Phage evolution

Exponentially growing bacteria and natural (ancestor) phage were mixed at final concentrations of approximately 2×10^6^ CFU/mL and 2×10^5^ PFU/mL (MOI 0.1), respectively, in a 96-well microplate with a total volume of 200 μL per well. Incubation followed at 37°C with shaking at 190 rpm for 24 h. Then, we added 0.1 volumes of chloroform to each well, and incubated for ten min with occasional mixing to terminate bacterial activity. We centrifuged the microplate at 4000 ×*g* for ten min to pellet the chloroform and bacterial debris. We then transferred the phage-containing supernatant to the corresponding well of a second microplate containing evolutionarily naïve exponentially growing bacteria. We repeated this process daily for up to ten days.

Bacterial growth kinetics curves (bacterial lysis by phage) with ancestral or evolved phages were obtained from sequential OD_600_ readings using a microtiter plate reader (Epoch Microplate spectrophotometer; BioTek, Winooski, VT, USA). We added bacterial cultures at exponential growth to individual wells of a 96-well microtiter plate and mixed them with phage lysate at an MOI of 1. The plate was incubated at 37°C in a microplate reader with continuous shaking and recorded the data every 15 mins for up to 24 h. Phage score was calculated using the bacterial areas under the growth curve (AUC) with and without phage^52,53^.

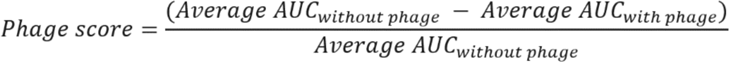

After choosing the focal day for each evolved phage, we purified a single phage genotype from the population by double single-plaque purification.

### Phage stability

To examine the stability of phages on long-term storage, we stored clean phage lysate in PBS in 500 µL volumes in 1.7 mL Axygen^®^ polypropylene microcentrifuge tubes (Corning Life Sciences, USA). These tubes were stored at ambient room temperature (∼22 °C), 37 °C and 4 °C. To track any change in stability, phage count was performed at the interval of 1, 3, 6 and 12 months, spanning up to 30 months.

### Bacterial and phage genomics and bioinformatics analysis

To extract bacterial DNA, we revived the bacteria from frozen stock and prepared overnight cultures. A 1.5 mL of overnight culture was used for DNA extraction using the GenElute™ Bacterial Genomic DNA Kit (Sigma-Aldrich, St. Louis, MI, United States) following the manufacturer’s protocol. For phages, we treated 1 mL of phage lysate with a concentration of >10^8^ PFU/mL with DNase (1 mg/mL) and RNase A (12.5 mg/mL) for 2 h at 37 °C, followed by an inactivation for 5 mins at 75 °C. Phage DNA extraction was performed using the Norgen phage DNA kit (Norgen Biotek Corp., Thorold, ON, Canada) following the manufacturer’s instructions. For cocktail’s (*Entelli-02*) genome, we extracted the genome from the final packed product following the same protocol mentioned above, excluding DNase and RNase treatment. We sent bacterial and phage DNA to Azenta Life Sciences in Suzhou, China for indexing and sequencing on an Illumina HiSeq platform.

The raw sequencing reads were trimmed using Trimmomatic v0.39^73^, followed by assembly using Unicycler v0.4^74^, annotation using Prokka v1.13.1^75^, NCBI PGAP^76^ and RAST annotation server^77^. Snippy v4.0 was used to identify nucleotide variation and SNPs. Phage genomes were assembled into complete genomes and aligned through pairwise sequence alignment using Pyani(https://huttonics.github.io/pyani/). Complete nucleotide sequences were rearranged to match the start gene position of the phage genome. Phages were assigned to the same genus if they had >95% Average Nucleotide Identity (ANI) with other known phages listed in the International Committee on Taxonomy of Viruses (ICTV) database. Phage genomes were examined against antibiotic resistance and virulence factor databases using the ABRicate tool (v1.0.1), and integrase genes in phage annotations were investigated. To analyse of prophage contamination in *Entelli-02,* prophage sequences on host genome were predicted using VirSorter2^78^ and verified manually to exclude falsely categorise or low-quality prophages with less than 5000 bp . The raw reads (fastq) of the *Entelli-02* were mapped with component phages’ genomes, respective production hosts and predicted prophages using Bowtie2^79^ and Samtools^80^ to obtain the ratios of read coverage and relative abundance (normalised coverage per million reads) as an indicator of prophage and host genome contamination.

### Complementation assay

We designed primers with suitable cut sites for the restriction enzymes targeting genes of interest (Supplementary Table 3). These primers were used to PCR-amplify the respective genes. Next, we performed two overnight double-digestion reactions at 37°C, employing the restriction enzymes, for each gene of interest and the vector pBBRMCS^69^. The digested products were purified through gel electrophoresis on a 0.7% agarose gel, followed by excision and ligation for 2 hours at room temperature using the Instant Sticky-end Ligase Master Mix (NEB) in accordance with the manufacturer’s protocol. The ligated product was then chemically transformed into *E. coli* DH5-α competent cells (NEB) following the company’s protocol. The transformed colonies were confirmed on antibiotic selection-LB plates and were subsequent validated through Sanger sequencing. The plasmid containing the desired complement was extracted from the confirmed colonies using GenElute™ Plasmid Miniprep Kit (Sigma-Aldrich). The plasmid containing the required complement was then electroporated into the phage-resistant mutants. To verify the restoration of phage infectivity, we conducted top agar and adsorption assays, with the anticipation of observing plaque formation and virion adsorption.

### *In vivo* assays (murine phage therapy)

We employed ECC isolate APO57, known to be permissive to all three phages in cocktail-V1, to establish a mouse infection model and determine the optimal bacterial density required to induce severe septicaemia. For the experiment, the bacterial inocula were prepared from an 18-hour culture, washed with PBS and adjusted to a concentration of 1×10^8^ CFU/mL (∼0.2 OD_600_). We used female 8-week-old BALB/c mice (18-20 g) for these experiments, with each mouse receiving either 1×10^6^ or 5×10^6^ bacterial cells, mixed with 6% porcine stomach mucin in PBS, for a total volume of 200 µL. Mice (*n* = 3 per group) were injected intraperitoneally and closely monitored for up to 12 h. When the humane endpoint or 12 hpi, whichever earlier, were reached, blood was collected via cardiac puncture, and a laparotomy was performed to obtain a liver section, right kidney, and spleen. The organs were weighed, homogenized in PBS, and plated to enumerate bacterial and phage counts, which were then normalised by organ weight or blood volume.

Following the establishment of the infection model, we proceeded with the phage treatment experiment. In this phase, each mouse group received a dose of 200 µL of 5×10^6^ CFU/mL. At 1 hpi the treatment group (*n* = 6) received 200 µL of PBS containing 1 × 10^8^ PFU/mL of phage, whereas the control group (*n* = 6) received an equivalent volume of sterile PBS. At the established endpoint, blood and organs were collected for bacterial and phage count.

### Phage production

Phage production was completed in two stages. In stage 1, each phage in the cocktail was amplified individually with its respective host in a 1000 mL batch. Here, 50 mL of an overnight culture of the host bacteria was added to 1000 mL LB broth supplemented with 1mM CaCl_2_ and MgCl_2_. This mixture was allowed to grow for ∼2 hours at 37°C with shaking until reaching an optical density of approximately 0.2 (OD600). At this point, phages were added to the mixture at an MOI of 0.1 and allowed to propagate overnight. Next, the lysate was clarified using sequential depth filters with retention rating of 0.5-5 µm (SUPRAcap 50 Pall^®^ Depth filtration capsule SCO50PDH4, Pall Corporation, Modultechnik GmbH, Germany) and of 0.2-3.5 µm (SCO50PDE2, Pall Corp) to remove bacterial aggregates. Pressure was monitored to ensure it did not exceed 2 bar to maintain filter specifications (with manufacturer’s maximum limit rating of 3 bar). The filtered lysate was then sterilized into sealed glass bottles within a biosafety cabinet through a sterilization grade filter with a 0.2 µm removal rating (Supor™ Mini Kleenpak™ capsules; KA02EKVP2G; Pall Corp) to ensure complete sterilization of the lysate. In stage 2, we purified and packaged the phages in a separate lab environment that was free from bacterial culturing. We used tangential flow filtration (TFF) with buffer exchange using ÄKTA Flux 6 system (product 29038438; Cytiva, Uppsala, Sweden) equipped with a 300 kDa nominal molecular weight cut-off (NMWC) microfiltration hollow fibre cartridge (Model; UFP-300-E-4MA, Cytiva). Briefly, the sterile lysate (∼900-1000 mL) was diluted up to 10-fold with 1× PBS and processed through TFF. This process was continued until the lysate lost its yellowish colour, indicating replacement of the bacterial media with the buffer, and was concentrated down to ∼150 mL final volume. The lysate was then collected and resterilised using a 0.2 µm filter as mentioned above. After clean-up and verification of the titre, the lysate of each phage was mixed, or diluted in 1× PBS as needed, to obtain a cocktail with a target final phage concentration of >10^8^ PFU/mL for each phage component. To clean endotoxins from the cocktail, we initially applied an EndoTrap^®^HD affinity column (Cat no. LET0035; Lionex GmbH, Germany) following the manufacturer’s instruction. However, the endotoxin level remained above the acceptable threshold. We therefore performed a 1-Octanol purification^71^. The endotoxin concentration at different stages of the purification process was assessed using the EndoZyme® II (Lot 22599; Hyglos GmbH (bioMérieux) Germany) and Endosafe® PTS™ cartridges (Product: PTS20F, Charles River Laboratories, MA, USA) kits. Once a satisfactory endotoxin level was achieved, the preparation was again filter sterilised and packed in 35 mL batches in sealed sterile glass vials (FILL-EASE™ Cat no. SV-50C02 Huayi Isotopes Company (Jiangsu, China) with syringe access. The vails were labelled and stored at 4 °C for clinical use. Third-party independent validation for sterility (US Pharmacopeia; USP 71) and endotoxin (USP 85) for each production batch was performed by Eurofins BioPharma Product Testing in Sydney, Australia. Phage titration was performed to verify the phage recovery throughout the process and product stability during storage.

### Statistical analysis

The statistical analysis was performed using R 4.2.3 and GraphPad Prism v9.1. Mean and standard error of mean, median, percentage and area under curve were calculated as appropriate, and statistical tests were performed accordingly and reported in figures where appropriate. Two-sided *p*-values were calculated and reported where applicable. The cut-off for the phage score was calculated using maximum likelihood estimation based on receiver operating characteristic curve implemented online^55^ (Supplementary Fig 14). Graphs were drawn using ggplot2 in R and Figures were drawn using Inkscape and Biorender.

### Ethical statement

All protocols involving animals were reviewed and approved by the Monash University Animal Ethics Committee (Project ID: 2022-27681-80733) and the animals were housed at the Monash Animal Research Facility, Monash University. We received all the bacterial isolates from the Department of Infectious Diseases at the Alfred Hospital without identifiable patient data. We conducted all procedures in strict adherence to institutional guidelines.

## Data availability (key resources table, lead contact, materials availability, data and code availability)

All data (supplementary data and figures) and results that were generated during this study are deposited at https://github.com/ECCphage.

Additional data are available in the Article, Online methods and Supplementary data/figure/table.

Genome sequences generated during this study are deposited at NCBI database under project ID PRJNA629076.

## Acknowledgements

The authors thank Professor Jian Li Monash University for providing plasmids used in this study, Associate Professor David McCarthy for raw sewage samples for phage isolation, researchers Leo Kan and Natasha Smith for their help in blind interpretation of spot assay results. We acknowledge Monash Animal Research Facility (MARP) for providing the necessary facilities for animal experiments; MASSIVE HPC facility (www.massive.org.au) for providing us with cluster time for data analysis and Monash Ramaciotti Cryo-EM platform for the use of facilities.

This work, including the efforts of Jeremy J. Barr, was funded by the National Health and Medical Research Council (NHMRC: 1156588 and 2026130).

## Author contributions

D.S, A.Y.P. and J.J.B. conceived the study. D.S, F.G.A. and A.P. performed phage isolation. D.S. performed phenotypic and genotypic characterization of phages, including all bioinformatics. D.S. and R.P. planned and performed complementation experiments. D.S. and R.D. performed phage evolution experiments. D.S. performed animal experiment, with help from F.G.A. (pilot study) and X.K. (pilot study). F.G.A. and L.B. performed spot assay and growth kinetics in the Alfred Hospital. N.M. provided strain information. D.K. performed TEM. J.J.B. and A.Y.P. supervised the project. J.J.B. funded the project. D.S. and J.J.B. wrote the original draft. All authors were involved in reviewing and editing the final manuscript.

